# Cell-surface targeting of fluorophores in *Drosophila* for rapid neuroanatomy visualization

**DOI:** 10.1101/2022.12.01.518740

**Authors:** Molly J. Kirk, Arya Gold, Ashvin Ravi, Gabriella R. Sterne, Kristin Scott, Evan W. Miller

**Affiliations:** Department of Molecular & Cell Biology, University of California, Berkeley, California 94720, USA; Department of Chemistry, University of California, Berkeley, California 94720, USA; Helen Wills Neuroscience Institute. University of California, Berkeley, California 94720, USA

## Abstract

Visualizing neuronal anatomy often requires labor-intensive immunohistochemistry on fixed and dissected brains. To facilitate rapid anatomical staining in live brains, we used genetically targeted membrane tethers that covalently link fluorescent dyes for *in vivo* neuronal labeling. We generated a series of extracellularly trafficked small molecule tethering proteins, HaloTag-CD4^1^ and SNAP_f_-CD4, which directly label transgene expressing cells with commercially available ligand substituted fluorescent dyes. We created stable transgenic *Drosophila* reporter lines which express extracellular HaloTag-CD4 and SNAP_f_-CD4 with LexA and Gal4 drivers. Expressing these enzymes in live *Drosophila* brains, we labeled the expression patterns of various Gal4 driver lines recapitulating histological staining in live brain tissue. Pan-neural expression of SNAP_f_-CD4 enabled registration of live brains to an existing template for anatomical comparisons. We predict that these extracellular platforms will not only become a valuable complement to existing anatomical methods but will also prove useful for future genetic targeting of other small molecule probes, drugs, and actuators.

## Introduction

Resolving the anatomical structure of the brain’s neural circuits is foundational to studying neuronal computations and behavior. The expression of genetically encoded fluorescent proteins is the most common method to explore neuroanatomy *in vivo*. Although expression of fluorescent proteins is a valuable technique, a limitation of the approach is that fluorescent proteins are often spectrally incompatible with other fluorophores and cannot be readily changed without the generation of different transgenic organisms.

For this reason, we sought to increase the flexibility of *in vivo* anatomical analysis. We proposed that an ideal system for *in vivo* anatomical analysis would 1) permit rapid and accurate staining of neuroanatomical structures *in vivo*, 2) facilitate changes in fluorescence spectra to readily pair with any available fluorophore, probe, or actuator, and 3) allow for temporal control of fluorescence addition and labeling. To increase the ease and flexibility of performing anatomical analysis *in vivo*, we have generated a series of extracellularly targeted small molecule tethering proteins that permit exploration of neuronal anatomy in a rapid, accurate, and highly flexible manner.

Although an *in vivo* approach offers a higher through-put method for anatomical analysis, the gold standard of anatomical techniques is immunohistochemistry (IHC). Here, tissues are preserved in a fixative and stained with antibodies targeted to specific proteins or epitopes.^2^ This technique allows for the multispectral labeling of multiple proteins and permits the highly detailed inspection of anatomical samples. However, this technique cannot be performed *in vivo*, as antibody access to intracellular epitopes requires permeabilization and fixation. This technique is often laborious, taking up to 2 weeks to achieve uniform staining in some preparations.^3^ A similar method, Hybrid IHC, shortens the timeline significantly.^3–5^ Hybrid IHC utilizes genetically encoded small molecule tethering systems, HaloTag,^6^ SNAP-Tag,^7^ and SNAPf,^8^ which traffic to the inner leaflet of the plasma membrane.^3^ These tethering platforms are genetically encoded monomeric enzymes that catalyze the formation of a stable covalent bond with ligand substituted fluorescent dyes.^9^ When the reactive dye is added to fixed and permeabilized tissues, it readily labels structures expressing the covalent tethering protein, mimicking the results of standard IHC. Hybrid IHC can label live murine skin samples *in vivo*.^10^ However, the intracellular location of the tethering proteins may limit dye binding in non-permeabilized tissues.

We sought to expand Hybrid IHC and develop an anatomical method which is genetically targeted, multispectral, and compatible with *in vivo* experimentation. To do this, we developed a series of extracellularly targeted small molecule tethering enzymes HaloTag-CD4^1^ and SNAP_f_-CD4, which covalently bind small molecules with the appropriate ligand (**Figure 1a)**.

**Figure 1.**
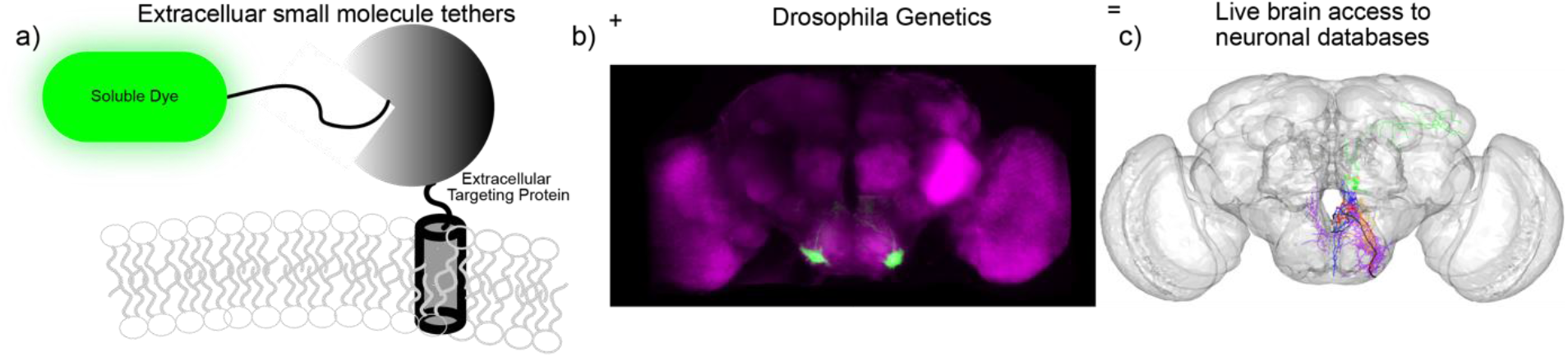
Chemical-genetic hybrids for neuron identification in live *Drosophila* brains. **a)** Commercially available Snap Tag and HaloTag reactive soluble dyes form covalent adducts with extracellularly targeted SNAP_f_-CD4 or HaloTag-CD4 molecules. **b)** The use of GAL4-UAS and LexA/LexA-op fly lines enables selective expression of SNAP_f_-CD4 and HaloTag-CD4 fusions in genetically defined populations of neurons in the fly brain. **c)** The extracellular location of these tethering enzymes permits the use of commercially available, water-soluble dyes across the visual spectrum and the registration of live brains to template space giving access from light level data to Gal4 expression pattern anatomical databases.

We then created stable transgenic *Drosophila* lines that express HaloTag-CD4 and SNAP_f_-CD4 on the extracellular surface in genetically defined neuronal populations (**Figure 1b**). Due to the extracellular localization of the tethering proteins, our method does not require permeabilization or fixation to access the covalent tethers and is thus amenable to the exploration of neuronal anatomy in live brain tissues (**Figure 1c**).

## Results and Discussion

### Generation of SNAPf constructs for expression in flies

Our group previously developed a chemical-genetic hybrid voltage sensor that targets HaloTag to the extracellular surface using an N-terminus PAT-3 secretion signal (from *C. Elegans*) and CD4 transmembrane anchor.^1^ We showed that HaloTag-CD4 could target dyes to the extracellular surface of neurons.^1^ We took a similar approach to generate a complementary extracellular labeling approach with SNAPf. We generated a PAT-3-SNAPf-HA-CD4 (SNAP_f_-CD4) fusion protein for extracellular trafficking: and sub-cloned it into both mammalian (pcDNA3.1) and insect expression vectors (pJFRC7, pJFCR19) **(Supplemental Scheme 1).**

SNAP_f_-CD4 shows good expression in HEK 293T cells: anti-CD4 immunocytochemistry confirms extracellular expression. The SNAPf self-labeling enzyme confirms not only localization but activity of the expressed enzyme by treating with Snap Tag reactive substrates. SNAP_f_-CD4 treated with SS-A488 (100 nM) shows good membrane localization in live HEK cells **(Figure 2a, Figure S1)**. Cells that express SNAP_f_-CD4 show approximately 6.5-fold higher fluorescence levels than non-expressing cells (**Figure 2b and c)**. Following live-cell imaging, cells can be fixed and retain their SS-A488 staining, which serves as a valuable counter stain to the anti-CD4 immunocytochemistry **(Figure S2).**

**Figure 2.**
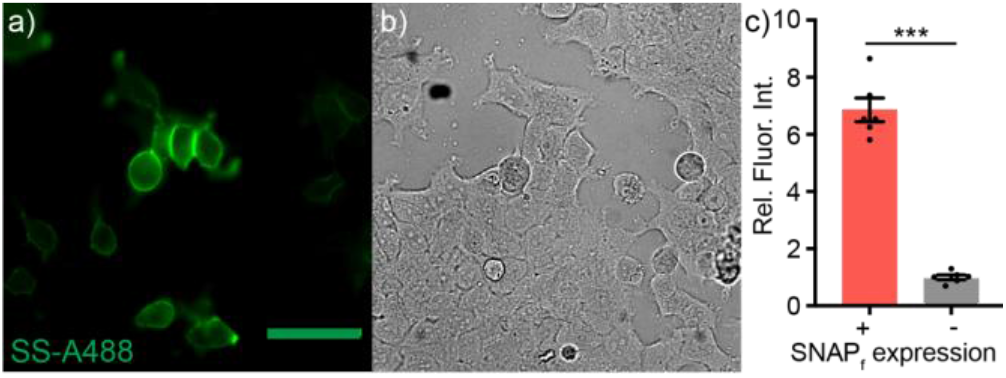
Live-cell staining with SS-A488 in HEK293T cells expressing SNAP_f_-CD4. Epifluorescence images of HEK293T cells expressing SNAP_f_-CD4 under the CMV promotor and **a)** stained with SS-A488 (100 nM, green). **b)** Transmitted light image of cells in panel **(a)** Scale bar is 20 μm. **c)** Plot of relative fluorescence intensity cells expressing and not expressing SNAP_f_-CD4. SNAPf (+) cells were assigned based on a threshold obtained from a non-transfected control. Data are mean ± SEM for n = 6 different coverslips of cells. Data points represent mean fluorescence intensity of 20-30 cells. (t-test, p< 0.0001)

We also observe cell surface localization of SNAP_f_-CD4 in S2 *Drosophila* cell lines, as visualized by anti-CD4 immunocytochemistry. S2 cells show SNAP_f_-CD4 dependent staining with SS-A488 (100 nM, **Figure 3, Figure S3**), with a 37-fold enhancement in fluorescence intensity in SNAP_f_-CD4 positive cells compared to non-expressing cells (**Figure 3c**). SS-A488 staining in S2 cells is also retained post-fixation **(Figure S4).**

**Figure 3.**
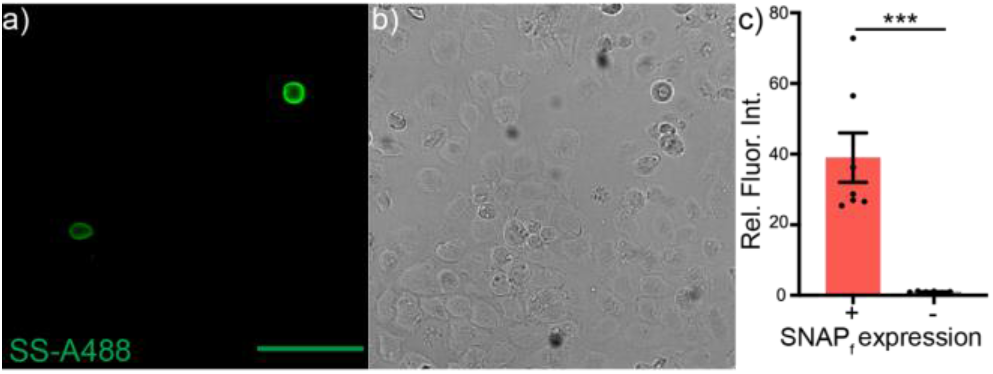
Live-cell staining in *Drosophila* S2 cells with SS-A488. Live-cell staining with SS-A488 in *Drosophila* S2 cells expressing SNAP_f_-CD4. Epifluorescence micrographs of S2 cells expressin SNAP_f_-CD4 under cotransfected pTubulin-Gal4 and **a)** treated with SS-A488 (100 nM). **b)** transmitted light image of panel **(a)**. Scale bar is 20 μm. **c)** Relative fluorescence intensities of SNAP_f_-CD4 expressing cells and cells that do not express SNAP_f_-CD4 from the same culture. SNAPf (+) cells were assigned based on a threshold obtained from a non-transfected control. Data are the ± SEM for n=7 cultures; data points represent the average fluorescence intensity of 20-30 cells. (t-test, p< 0.0001)

### Validation of UAS-SNAP_f_-CD4 transgenic fly lines

To evaluate the performance of cell surface-expressed SNAP_f_-CD4 in live brains, we generated transgenic flies for both Gal4/UAS (pJFRC7) and LexA/op (pJFRC19) **(Figure S5)** expression (Best Gene Inc.). Crossing the resulting UAS-SNAP_f_-CD4 line with a pan-neuronal driver line, neuronal synaptobrevin-GAL4 (nSyb-GAL4) results in SNAPf expression in all neurons in the *Drosophila* brain.^11^ Brains of nSyb-GAL4>SNAPf-CD4 flies show robust CD4 and HA expression **(Figure 4a, Figure S5a and d)**, unlike controls which do not express SNAP_f_-CD4 or the HA epitope tag **(Figure 4c, Figure S5b and e)**. The anti-CD4 and anti-HA fluorescence pattern indicates good localization to the plasma membrane, as the immunofluorescence is isolated from the Hoechst nuclear counterstain in confocal optical sections **(Figure 4b).**

**Figure 4.**
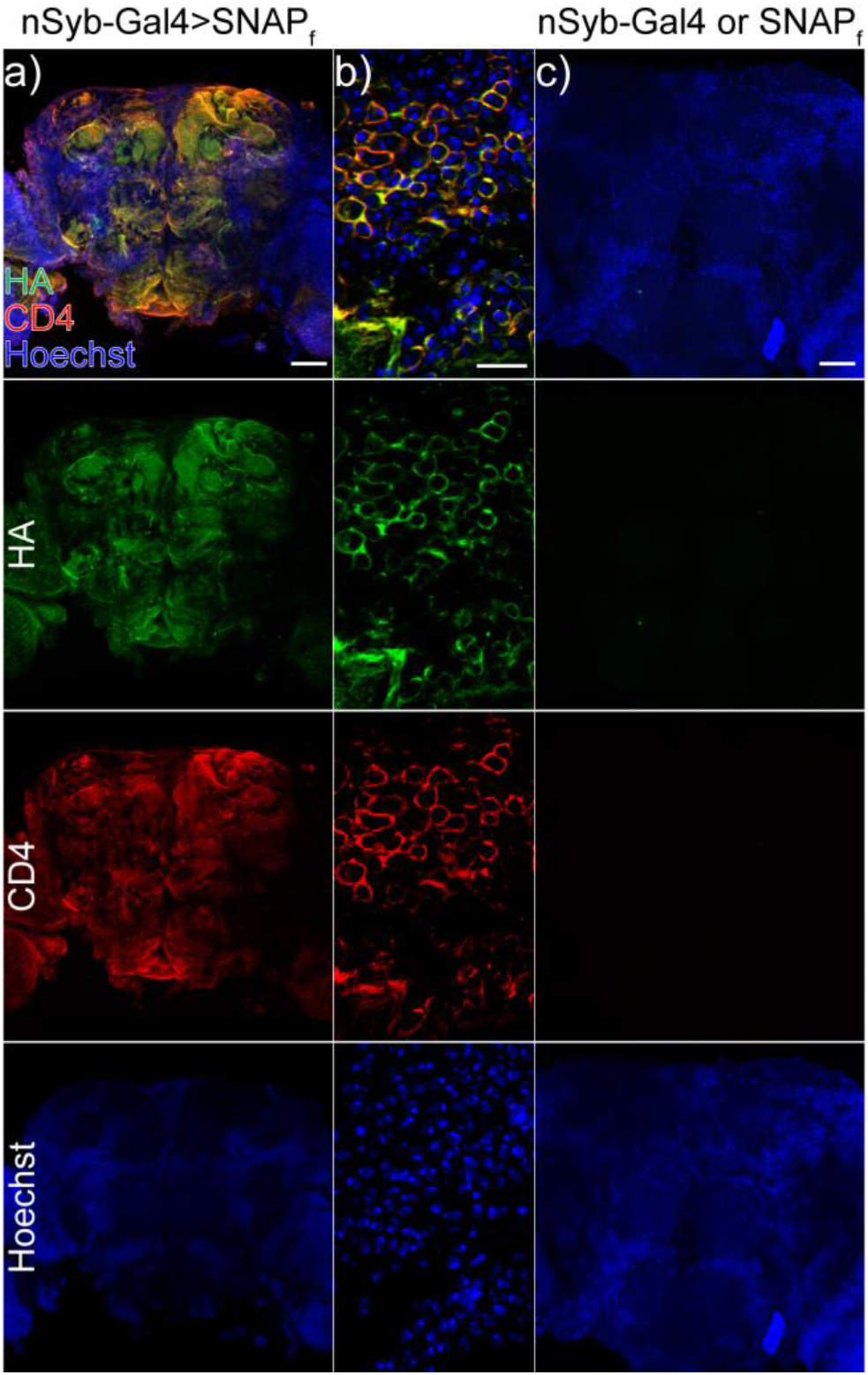
Immunohistochemistry of pan-neuronal SNAP_f_-CD4 expression and trafficking in *Drosophila* brain. **a)** Maximum confocal z-projection of a fixed nSyb-Gal4, SNAP_f_-CD4 brain stained under non-permeabilizing conditions for HA (green), CD4 (red) and nuclear counterstained using Hoechst 33342 (at a concentration of 10 μg/μL, equivalent to 16 μM) **b)** 63x single confocal plane of cells expressing SNAP_f_-CD4 under the nSyb-Gal4 driver line as shown in panel **(a)**. **c)** maximum confocal z-projection of either nSyb-Gal4 or SNAP_f_-CD4 brain, which does not express the SNAP_f_-CD4 protein. Scale for all images is 50 μm.

### Development and assessment of live brain Hybrid IHC

In developing our dye loading protocol, we aimed to limit time required to perform the technique and number of tissue interactions. In **Figure 5a,** we schematize IHC and Hybrid IHC, highlighting each method in terms of these two as-pects. IHC takes up to 12 days and requires over 15 tissue interactions. Hybrid IHC dramatically decreases this time to approximately 1 hour with 8 tissue interactions. SNAP_f_-CD4 and HaloTag-CD4 extracellular tether dye loading protocol requires only 1-2 tissue interactions and takes approximately 15 minutes in total. The short labeling time with SNAP_f_-CD4 and HaloTag-CD4 may facilitate assessment of neuronal anatomy after functional imaging studies.

**Figure 5.**
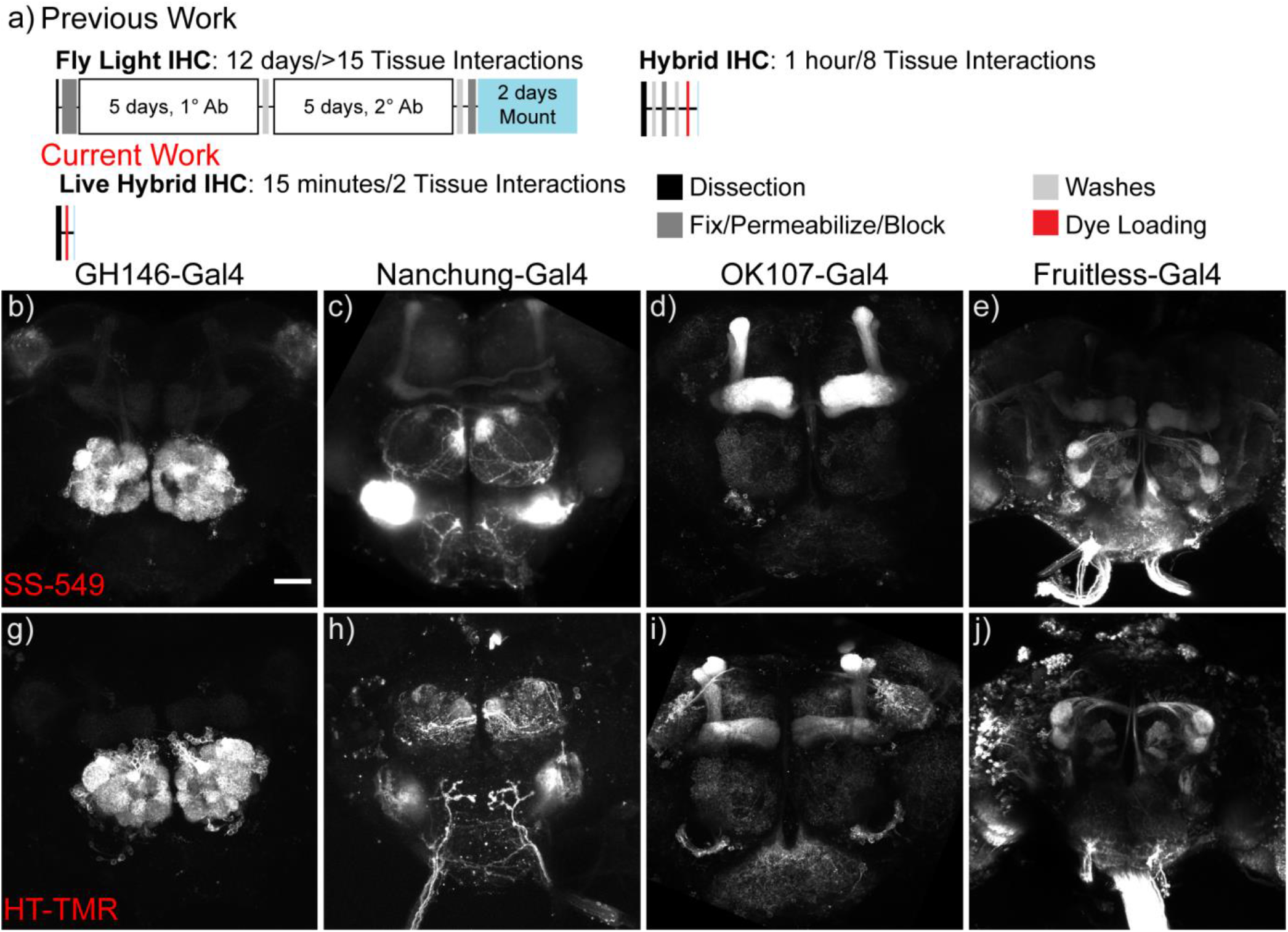
Dye loading of Gal4 expression patterns in explant brains. **a)** timeline comparing time required for common immunohistochemistry protocol from Fly Light (top), Hybrid IHC (middle), and live Hybrid IHC (bottom, the method reported in this manuscript). Timeline includes total time required for each technique and the breakdown of each technical step and number of tissue interactions (TI). Diagram is not to scale. Maximum confocal z-projection of live SS-549 dye (1 μM) loading in *Drosophila* explant brains expressing SNAP_f_-CD4 under **b)** GH146-Gal4, **c)** Nanchung-Gal4, **d)** OK107-Gal4, and **e)** Fruitless-Gal4. Maximum confocal z-projection of live HT-TMR dye (1 μM) loading in explant brains expressing HaloTag-CD4 under **g)** GH146-Gal4, **h)** Nanchung-Gal4, **i)** OK107-Gal4, and **j)** Fruitless-Gal4. Scale for all images is 50 μm.

We expressed SNAP_f_-CD4 and HaloTag-CD4 in fly brains by crossing with four commonly used Gal4 driver lines, GH146-Gal4 **(Figure 5b),** Nanchung-Gal4 **(Figure 5c)**, OK107-Gal4 **(Figure 5d),** and Fruitless-Gal4 **(Figure 5e)**. To evaluate the ability of SNAP_f_-CD4 and HaloTag-CD4 to label a variety of Gal4 expression patterns with varying depth, complexity, and specificity, we loaded these live brains with either HaloTag-Tetramethyl Rhodamine (HT-TMR)^1^ or Surface Snap-549 (SS-549, 1 μM NEB). We found that these dyes robustly labeled the expected neuronal populations for each line regardless of the number of neurons or their location within the brain and showed high-intensity staining with minimal background fluorescence.

SNAP_f_-CD4 labeling of genetically defined cell populations is not only fast, but flexible, since it is compatible with a wide range of spectrally-tuned dyes **(Figure 6a).** To illustrate this, we loaded GH146-Gal4>SNAP_f_-CD4 brains with SS-A488, a green dye **(Figure 6b)**, SS-549, a red-shifted dye **(Figure 6c)**, or SS-A647, a far red-shifted dye **(Figure 6d)**. These live brains show the expected expression pattern in the antennal lobe, labeling both cell bodies and dendritic fields with high intensity and minimal background staining. The “plug and play” feature of SNAP_f_-CD4 and HaloTag-CD4, i.e., the ability to rapidly switch colors to suit the needs of a specific experiment, distinguishes our approach from the expression of fluorescent proteins. We generated the LexA/lexAop^12^ versions of both SNAP_f_-CD4 and HaloTag - CD4 to showcase the genetic as well as chemical versatility of this approach (**Figure S6**).

**Figure 6.**
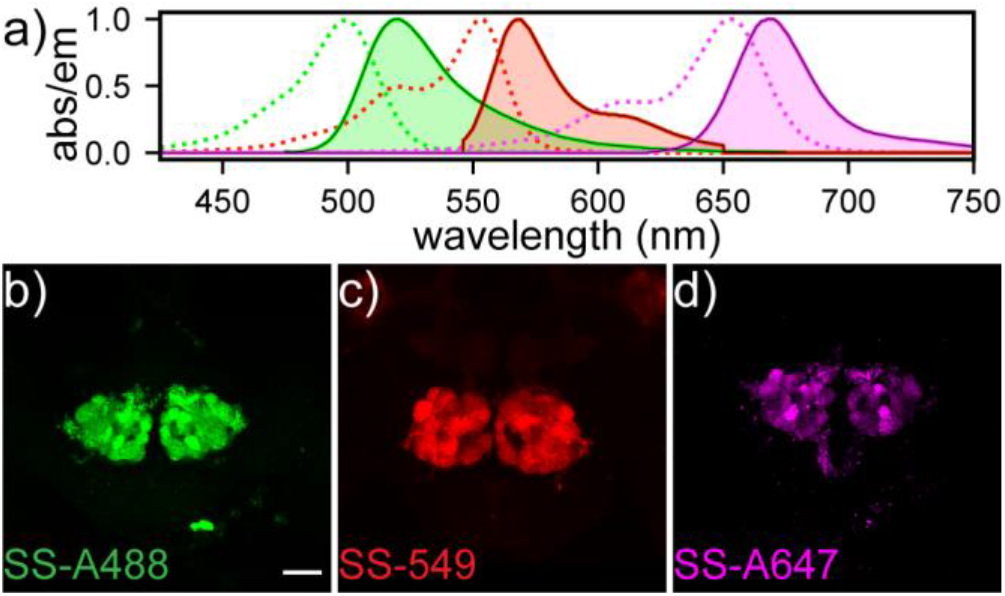
Dye loading in *Drosophila* explant brains. a) excitation (dotted lines) and emission (solid lines) spectra for commercially available SS-A488 (green), SS-549 (red), SS-A647 (magenta). Maximum z-projections of GH146-Gal4, SNAP_f_-CD4 live fly brain explants brains stained with **b)** SS-A488, **c)** SS-549 and **d)** SS-A647 (1 μM, all dyes). Scale is 50 μm.

### Live brain registration using SNAP_f_-CD4

We next sought to register live brains to an anatomical template brain to allow for direct comparisons of anatomy across different specimens. During the registration process, brain images are transformed via rigid and non-rigid transformations to match the coordinate space of an anatomical template brain.^13,14^ Once transformed to template space, one can directly compare the expression patterns and single-cell projection patterns to existing anatomical data-bases such as FlyCircuit^15^ using the similarity algorithm NBLAST.^16^ Registration is most often performed using immunostaining for BRP (bruchpilot),^17^ a pan-neuronal marker for neuropil regions, which creates a strong counterstain for neuronal anatomy. Here, the BRP staining must be uniform and permeate evenly throughout the sample for the registration to properly align the samples.

Uniform staining throughout the brain is a prerequisite for registration. Fixed and permeabilized brains expressing an intracellular SNAPf::BRP fusion protein and stained under permeabilizing Hybrid IHC conditions show uniform staining.^4^ However, when we applied those same SNAPf::BRP brains to our live brain staining protocol (using cell-permeable dye JF-546),^18^ we observe fluorescence mainly in superficial layers of the live brain and uneven signal in deep brain structures (**Figure S7**). Pan-neuronally expressed nSyb-Gal4>SNAP_f_-CD4 loaded with cell impermeant SS-A647 showed uniform and penetrant staining.

Pan-neuronal SNAP_f_-CD4 labeled with SS-A647 (**Figure 7a**) closely recapitulates BRP immunohistochemistry of the JFRC2010 template brain,^14^ which was generated from a single female brain immunostained for BRP (**Figure 7b**). We thus hypothesized that the pan-neuronal SNAP_f_-CD4 could be used to register brains to a template space directly. Testing this hypothesis, we loaded nSyb-Gal4>SNAP_f_-CD4 with SS-A467(10 μM) and found that we could readily register the first half of the brain (~100 μm in depth) to the JFRC2010^14^ template using the CMTK registration algorithm.^19^ To assess the efficacy and accuracy of our staining, we generated a data set of 20 female and 20 male live-loaded brains, which we registered to JFRC2010 **(Figure 7a-c).** We determined successful registrations by overlaying the registered live brain data **(Figure 7a)** with the template brain **(Figure 7b)** and qualitatively comparing the regional overlap of the various structures within the brain **(Figure 7c)**. We had a50% success rate across all brains sampled. Brains that failed to register often had poor alignment due to a poor imaging orientation or minor damage to tissue resulting from the dissection process.

**Figure 7.**
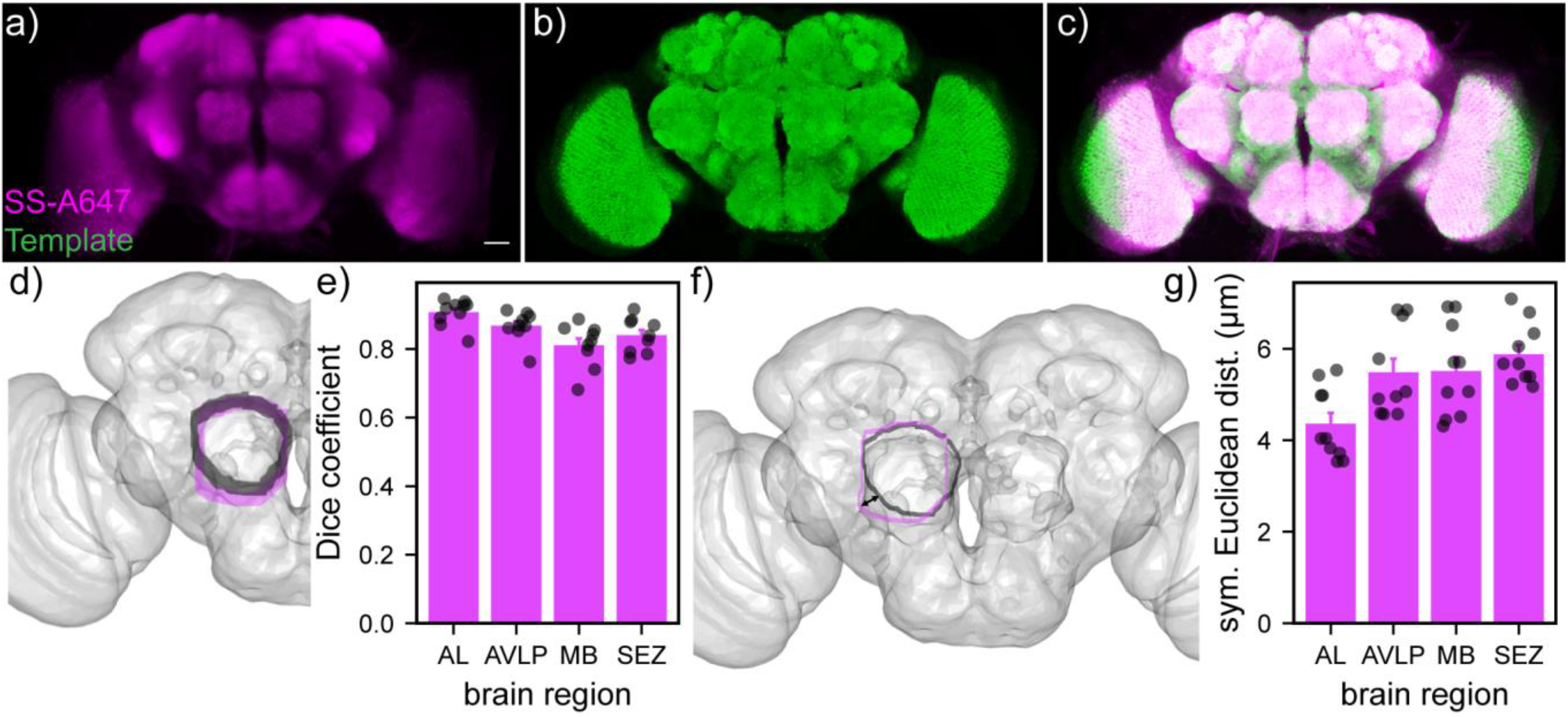
Live brain registration using SS-A647 and quantification of registration quality. Explant nSyb-Gal4>SNAP_f_-CD4 brains were loaded with SS-A647 (10 μM) then registered to JFRC2010 template space using CMTK registration algorithm. **a)** Average z-projection of mean live brain data, constructed from the pixel-wise average of 10 individual confocal stacks of live brains stained with SS-A647 (5 male and 5 female) all registered to JFRC2010. **b)** Average z-projection of JFRC2010 template brain confocal image. **c)** 3D rendering of merged template (green) and mean live brain data (magenta). Scale for all images is 50 μm. **d)** Visual schematic of Dice Coefficient measure of areal overlap, where area of the Ito ROI (Grey) is compared to the area of the experimenter drawn ROI (magenta) using the equation depicted below. R^T^ is the area of the Ito ROI, and RD is the area of the drawn ROI. **e)** Average Dice Coefficient ± SEM of the best-registered slice from each region. Each data point represents the Dice Coefficient from one slice of an individual brain registered to JFRC2010 (n= 10, 5 male, 5 female). **f)** Schematic of the measurement obtained by Symmetric Euclidean Distance where the shortest distance from one ROI to another is averaged (dD is shortest distance from Ito ROI to the drawn ROI and d^T^ is the shortest distance from the drawn ROI to the Ito ROI). **g)** Plot of average Symmetric Euclidean Distance for each individual brain (n=10, 5 female and 5 male) across all z planes for a specific region.

We quantitatively evaluated the registration using two independent measures: Dice coefficient and mean symmetric Euclidean distance.^20^ Dice coefficient measures the areal overlap between a registered brain region and its corresponding region^20^ on the template brain and is defined by Equation 1:

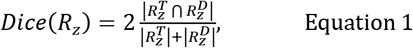

Where R^T^ is the area of a template-defined ROI^21^ and R^D^ is the user-defined region of interest (ROI). The Dice coefficient has an upper bound of 1 (complete overlap between the template and user ROIs) and a lower bound of 0 (no overlap).

We compared ROIs throughout each brain structure in 5 μm steps throughout the volume **(Figure 7d, Figure S8)**. We chose four brain regions, antennal lobe (AL), anteroven-trolateral protocerebrum (AVLP), mushroom body (MB), and suboesophageal zone (SEZ), which vary in both depth within the brain and anatomical structure. All four brain re-gions show high Dice coefficients at their best-registered z-plane, ranging from 0.9 (AL) to 0.8 (MB), indicating a high level of overlap with the template ROI **(Figure 7e, Figure S8).**

As another measure of the goodness of fit of our live brain registration approach, we calculated the average boundary error as the mean symmetric Euclidean distance (SED) or the average of the shortest distance between the template ROI border and the experimenter drawn ROI border and the symmetric computation **(Figure 7f),** ^20^ defined by Equation 2:

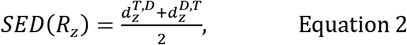

Where d^T,D^ is the mean distance between each point on the border of the template ROI and the closest point on the user-defined ROI, and d^D,T^ is calculated by determining the symmetric relationship.

Using this method, we find that the average boundary error is between 4-6 μm across all four brain regions **(Figure 7g)**. This resolution is 10× larger than reported for immunostaining registration to template, which can achieve resolution of ~ 0.4 μm.^20^ The average boundary error (SED) could be improved by generating a live template brain that can be bridging registered to the JFRC2010 template space. We calculated the error for each z-plane at 5 μm steps within the volume. We find that central portions of most brain regions show low boundary error (**Figure S9**); the extremes of structures tended to show the most dramatic boundary error. This trend also appears in the Dice coefficient **(Figure S8)**.

To further assess the quality of registration using live staining and access anatomical databases via similarity searches, we performed an NBLAST search using live registered brains. We expressed GFP::CD8 under a sparse LexA driver line R34G02-LexA, which labels a single bilateral neuron pair in the suboesophageal zone called interoceptive neurons or ISNs.^22^ We registered these brains using pan-neuronally expressed SNAP_f_-CD4 labeled with SS-A647. Manually tracing the right projection pattern of the ISN produced the target neuron for our search (**Figure 8a**). To create our query database, we seeded a database of 15,500 single neurons^15^ and expression patterns^23^ with a manually traced ISN neuron from an immunostained and registered brain (**Figure 8b)**. We then performed an NBLAST search using the live registered brain and the seeded database. Less than 0.2% all neurons returned a positive NBLAST score (32 out of 15,501, **Figure 8e-f**). The top 5 hits of the search (**Figure 8d,e**) all have neuronal arbors that overlap with the live-stained ISN query neuron (**Figure 8d**, red) and the immunostained ISN standard (**Figure 8d**, black).

**Figure 8.**
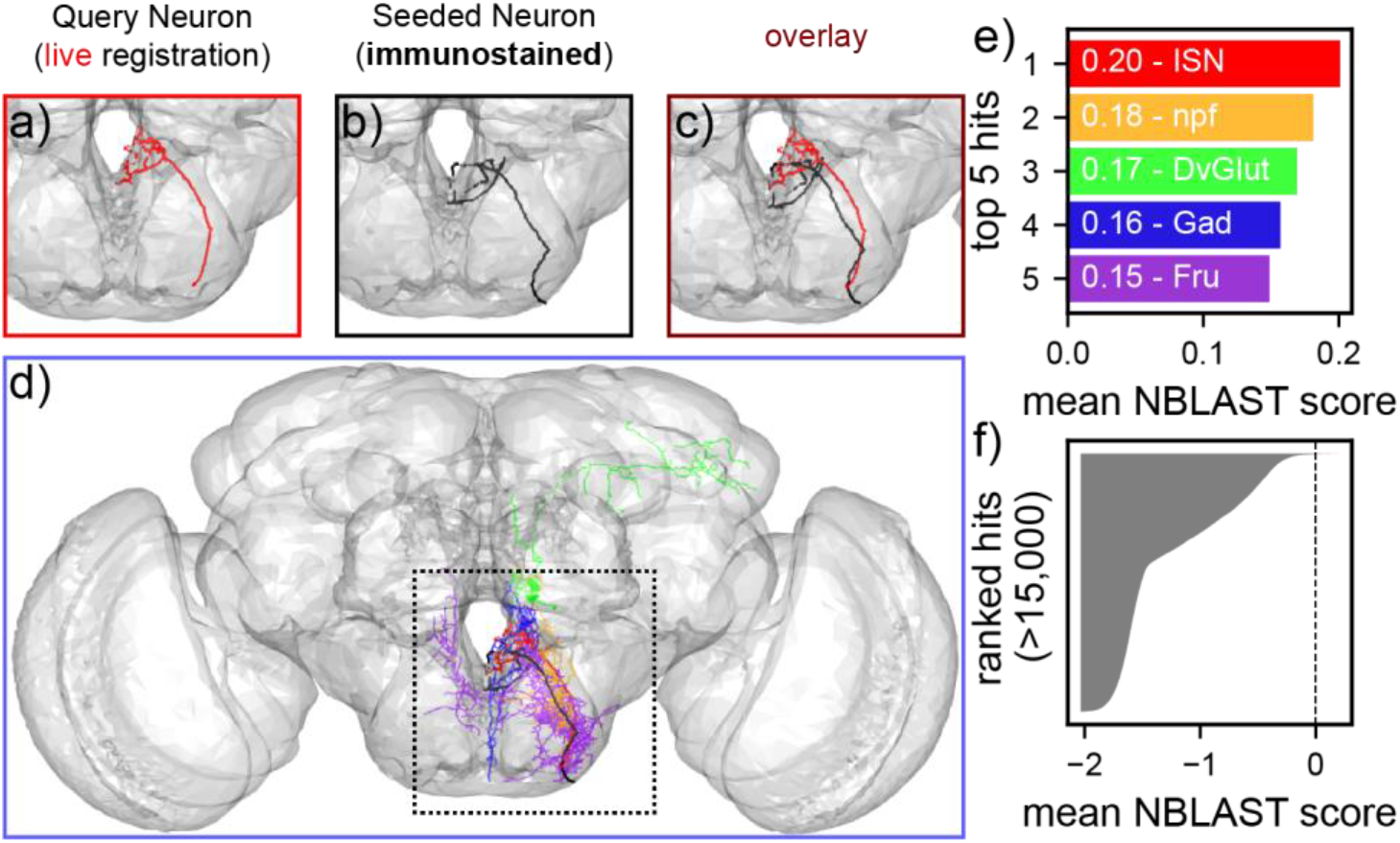
NBLAST search using live registered brain samples. The target ISN neuron manually traced expression pattern of R34G02-LexA>CD8::GFP and registered to JFRC2 using SNAP_f_-CD4 tethered SS-A647 **(a,** red). Data was further bridged to a shape-averaged template brain (FCWB, see Chiang, *et al. Curr. Biol*. **2011**, *21*, 1-11) for analysis. **b)** Query database seed for the ISN neuron generated from fixed and immunostained ISN neuron sample registered to JFRC2 and bridged to FCWB (black). **c)** Overlay of live brain ISN neuron (red) and fixed brain ISN neuron (red) registered to JFRC2 and bridged to FCWB. **d)** Overlay of top 5 hits from NBLAST query. Dotted line shows regions in panels a-c. **e)** Mean NBLAST scores of the top 5 hits. Full names are in Table 1. **f)** Mean NBLAST scores of all 15,500 queried neurons.

**Table 1.**
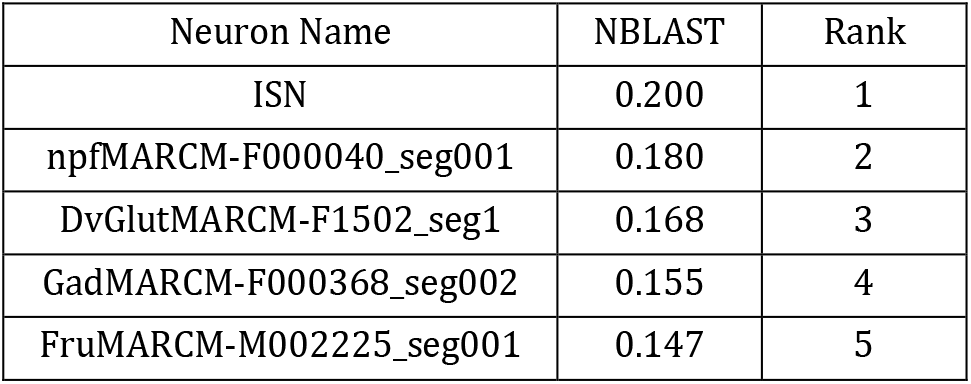
Top hits of NBLAST search.

The immunostained ISN tracing was the top hit out of 15,500 neurons, with a mean normalized NBLAST score of 0.2 (**Figure 8e-f**).

## Conclusion

In summary, we show that extracellularly anchored small molecule tethers HaloTag-CD4 and SNAP_f_-CD4 can be used to covalently label genetically defined populations of cells with commercially available dyes both in cultured cells and in live *Drosophila* brain tissue. We show that UAS-SNAP_f_-CD4 and LexA-SNAP_f_-CD4 (**Figure S6)** can be used to rapidly stain and label brains (15 minutes vs 1 hour or 12 days) and then directly register brains to template space with up to 90% overlap between live-brain determined ROIs and template and ~5 μm boundary error. We use live registered brains to access neuronal morphology databases using the similarity search NBLAST, showing that the live-brain registration approach can be readily incorporated into existing workflows. The hybrid chemical-genetic nature of our system provides a high level of flexibility: a broad range of dyes are available for immediate application and may be selected for use without the generation of a novel transgenic fly. This multiplexing of anatomical and physiological experimentation can be used simultaneously or sequentially within the same live sample.

Despite these advances, several drawbacks are associated with our current method and represent opportunities for improvement. First, our approach is entirely Gal4/LexA dependent and thus may not yield satisfactory results in weakly expressing lines. This issue may be addressed via the addition of tandem repeating small molecule targeting proteins, which can be secreted to the extracellular surface. Alternatively, the expression of UAS-Gal4 or lexAop-LexA reporter lines to increase the expression of the activator protein in positive cells, increasing staining efficiency.

Second, we are only able to register the first half of the brain. We hypothesize this is a consequence of poor penetration of the small molecule fluorophores through the brain. Rhodamine dyes, with high two-photon cross sections^24^ and improved solubility^25^ may help.

Finally, our current methodology is not as accurate at registering brains or revealing fine neuronal arborizations as IHC^20^ or Hybrid IHC^26^ as shown by the larger boundary error and inability to visualize small neurites in the GFP labeled expression patterns, and low, yet non-zero NBLAST scores. In specific situations, these issues may hinder its usefulness in neuronal identification. This is potentially due to the need for rapid acquisition (under 15 minutes) of images to prevent morphological changes to the brain structure in explant brains and could be mitigated by the application and imaging of the brain in the intact fly. Future endeavors should focus on optimizing the system for high-resolution microscopy in the intact fly and determining best practices for registration, including algorithm, sample preparation, and template selection.

It is important to note that our study is not the first to register live brains to a template. Other groups have been able to do this by registering intracellularly expressed Td-Tomato imaged under two-photon excitation.^27–29^ Although these tools can be used to register whole brain, they have not been used to perform morphology similarity searches *in vivo*. Our technique complements existing systems by offering spectral flexibility and rapid visualization. These tools may be adopted for live brain staining as presented here and will also be useful for genetic targeting of other small molecules such as drugs, actuators, and indicators to the extracellular surface of genetically defined cells in the fly brain.

## ASSOCIATED CONTENT

Experimental methods and details, supporting figures.

## ACKNOWLEDGMENT

We acknowledge support from NIH (R01NS098088, EWM) and NSF (NeuroNex Innovation Award 1707350, EWM and KS). MJK was supported, in part, by a training grant from the NIH (T32GM007232.

