## Supporting Information Document for "Cell-surface targeting of fluorophores in *Drosophila* for rapid neuroanatomy visualization"

**Affiliations**:

*Corresponding Author

**DOI**: Placeholder

**Figure S9.** Dye loading in panneuronal LexA lines .......... 15

**References.** .......... 19

**Methods:**

**Plasmid construction.**

We included a secretion signal derived from the signal peptide of the *Caenorhabditis elegans* β-integrin PAT-3 at the N-terminus of $SNAP_f$, with the 5' UTR from heat shock protein 70 (hsp70) and the 3'UTR and poly A tail from SV40 early genes, as described previously.[1] For expression in HEK 293T cells, we subcloned $SNAP_f$ via restriction digest (NheI, SalI) and subsequent Gibson Assembly into pCDNA3.1(Thermo Fisher) vector containing a cytomegalovirus (CMV) promoter, a 5' PAT3 secretion signal, and a 3' CD4 transmembrane domain. For expression in S2 cells and transgenic generation, the insert Pat3-SNAPf-HA-CD4 was assembled into pJFRC7[2] backbone via restriction digest (XhoI and XbaI removed CD8::GFP) and Gibson assembly (Addgene). All constructs were sequence confirmed by the UCB Sequencing Facility. Sequences used for all constructs can be found in the attached electronic construct maps.

**Scheme S1.** Construct maps of mammalian and *Drosophila* constructs

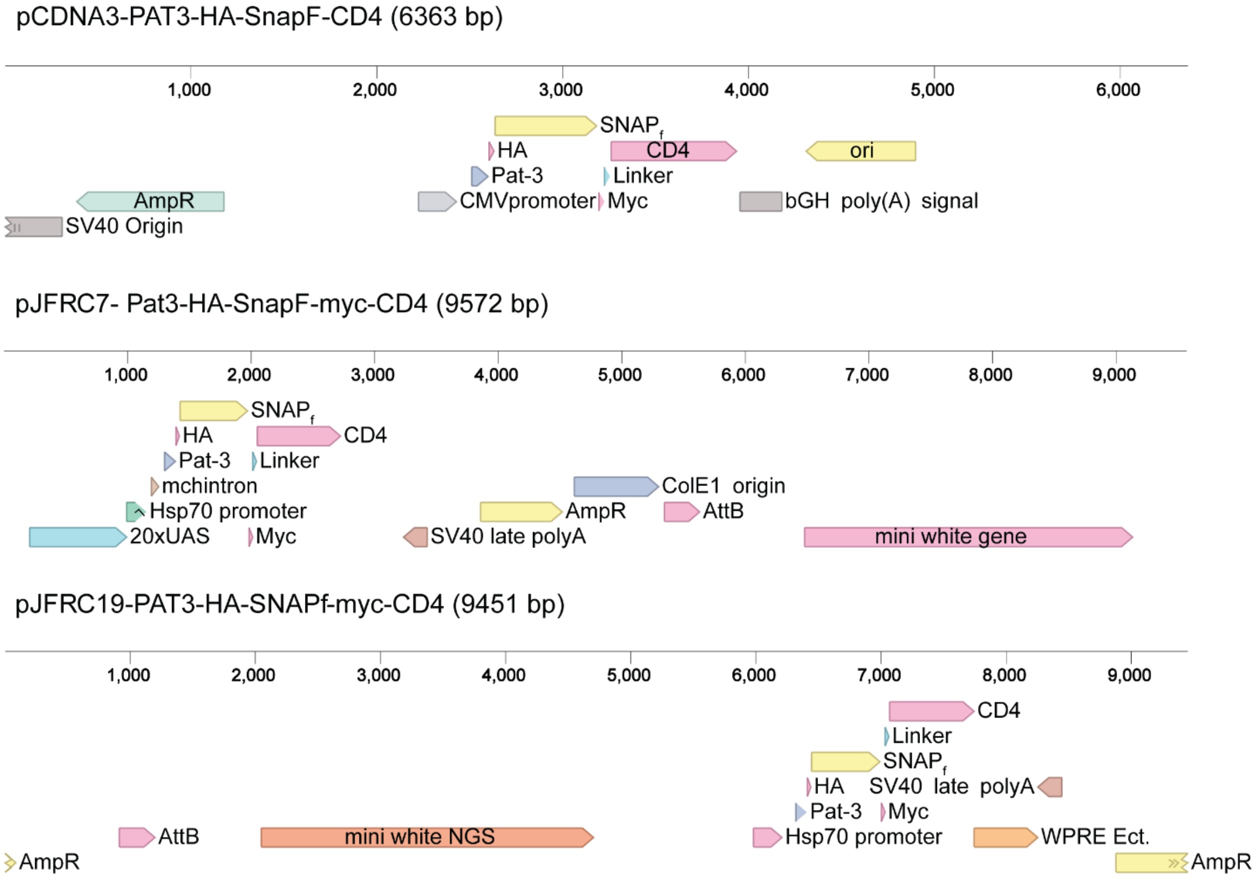

**Cell culture and transfection.**

We obtained all cell lines from the UCB Cell Culture Facility. Human embryonic kidney 293T (HEK) cells were maintained in Dulbecco's modified eagle medium (DMEM) supplemented with 1 g/L D-glucose, 10% fetal bovine serum (FBS; Thermo Scientific), and 1% GlutaMax (Invitrogen) at 37 °C in a humidified incubator with 5 % CO2. Cells were passaged and plated in DMEM (as above) at a density of 50,000 cells onto 12 mm coverslips pre-treated with Poly-D-lysine (PDL;1mg/ml; Sigma-Aldrich). Plasmid transfection was carried out using Lipofectamine 3000 (Invitrogen) 12 hours after plating. Imaging was performed 36 hours after plating.

S2 Cells were maintained in Schneider's *Drosophila* media (Thermo Fisher Scientific) supplemented with 10% FBS at 28°C in a non-humidified incubator under atmospheric conditions. Cells were passaged and plated at 500,000 cells per well in a 24 well plate. Six hours after plating, Tubulin Gal4 pCaSper (Addgene #17747) was cotransfected with pJFRC7 constructs using a modified Lipofectamine 3000 (Life Technologies) protocol. This protocol included a 20-minute preincubation of lipofectamine and DNA in Opti-MEM (Life Technologies) and no p3000 reagent. Forty-eight hours after transfection, S2 cells were transferred onto PDL (1 mg/mL) treated 12 mm coverslips and allowed to adhere for 30 minutes before dye loading and imaging.

**Dye loading.**

We maintained DMSO stock solutions (100 μM) of all dyes at -80 °C in single-use aliquots. Aliquots were further diluted to a working concentration of 100nM in HBSS and incubated with cells for 30 minutes at 37 °C for HEK cells and room temperature for S2 cells. We then replaced all dye-containing HBSS with fresh HBSS and imaged in HBSS at room temperature.

**Epifluorescence microscopy.**

Imaging was performed on an AxioExaminer Z-1 (Zeiss) equipped with a Spectra-X Light engine LED light (Lumencor), controlled with Slidebook (v6, Intelligent Imaging Innovations). Images were acquired with a W-Plan-Apo 20x/1.0 water objective (20x; Zeiss) and focused onto an OrcaFlash4.0 sCMOS camera (sCMOS; Hamamatsu). The optical setup for imaging with each dye is described below.

**Table S1.** Optical filter sets for epifluorescence microscopy

| Dye | Excitation | Emission | Dichroic |
|---|---|---|---|
| mSNAP2, SS-A488 | 475/34 nm BP | 540/50 nm BP | 510 nm LP |
| A647 | 542/33 nm BP | 650/60 BP | 594 nm LP |
| Hoechst 33342 | 375-400nm | 405/40 BP | 415 LP |

**Epifluorescent image analysis.**

For fluorescence intensity measurements, regions of interest were drawn around cell bodies, and fluorescence was calculated in ImageJ (FIJI), NIH). We identified transfected cells by setting a threshold that excluded all cells in the non-transfected controls. We calculated the fold change between non-transfected and transfected cells by taking the ratio of transfected cells fluorescence and non-transfected cell fluorescence. For each condition, at least 100 cells were circled across three to five individual coverslips.

**Immunocytochemistry.**

Immediately following live-cell dye loading experiments, cells were fixed for 20 minutes at room temperature with 4% formaldehyde in PBS. Cells were washed in PBS (3x 5-minute washes) and treated with either 0.3% Triton X-100 in PBS for the permeabilized condition or PBS for the non-permeabilized condition. Cells were again washed in

PBS (3x 5-minute washes) and blocked for at least 45 minutes in 0.1% NGS in PBS. Cells were then incubated overnight at 4 °C with 1:500 mouse anti CD4 (OKT4; Thermo Fisher Scientific). We then washed each sample in PBS and stained with a spectrally compatible Goat anti-Mouse A594 (Life Technologies) 1:1000 in 0.1%NGS for 2hrs at room temperature. We added Hoechst 33342 (10mg/mL, -20 stock) 1:1000 for the last 15 minutes of this incubation period. Cells were washed (3 x 5-minute washes) in PBS and mounted onto glass slides using Fluoromount Mounting Media (VWR International) before imaging.

**Transgenic generation.**

pJFRC7-Pat3-SNAP$_f$-CD4 were sent to Best Gene Inc. for injection into the following genomic sites via phi-C31 integration.

***Table S2.*** *Injection phi C31 site and stock line*

| Construct | Injection Site | Injection Stock |
|---|---|---|
| pJFRC7-Pat3-SNAP$_F$-CD4 | VK05 | #9725 |

**Immunohistochemistry.**

Fly brains were dissected in calcium magnesium-free artificial hemolymph (AHL-/-; NaCl 108.0 mM, KCl 5.0 mM, $NaHCO_3$ 4.0 mM, $NaH_2PO_4 \cdot H_2O$ 1.0 mM, Trehalose· 2 $H_2O$ 5.0 mM, Sucrose 10.0 mM, HEPES 5.0 mM and adjusted to pH 7.5 with NaOH) and fixed for 20 minutes in 4% formaldehyde in PBS. Brains were then washed in PBS (3x 5-minute washes) and treated with either 0.3% Triton x -100 in PBS for the permeabilized condition or PBS for the non-permeabilized condition. Brains were again washed in PBS and blocked for at least 45 minutes in 0.1% NGS in PBS. Cells were incubated for 5 days in block containing 1:50 RT anti HA (Sigma Aldrich) and 1:100 mouse anti CD4 at 4 °C. Brains were then washed in PBS (3 x 5-minute washes) and stained with Goat anti-Rat A488 (Life Technologies) 1:1000 and Goat anti-Mouse A594 (Life Technologies) in block for 4 hours at room temperature shaking. We added Hoechst 33342 (10 mg/mL) 1:1000 for the last 15 minutes of this incubation. Brains were then washed and mounted onto glass slides using Vectashield mounting media (Vector Laboratories) before imaging using confocal microscopy.

**PLL Coverslips.**

PLL was prepared using the FlyLight recipe. (https://www.janelia.org/sites/default/files/FL%20Recipe%20-%20Poly-L-Lysine%20.pdf). Briefly, PLL (Sigma Aldrich. # P1524-25MG) was thawed to room temperature and diluted with 2mL of double-distilled water, transferred to a 50 ml conical vial, and further diluted with 30mL of double-distilled water. 64uL Photo-Flo (Electron Microscopy Sciences. # 74257) was added to this mixture and vortexed. Coverslips were then dipped into PLL and placed on a Kim wipe while drying overnight. Once dried, coverslips were stored at -20 until use.

*Dye loading and imaging for expression patterns*

Flies aged 10 days post eclosion were dissected in ice-cold AHL-/-. Brains were stored on ice in calcium magnesium-free AHL during the remainder of the dissections (no longer than 20 minutes) before loading with 1 µM Surface Snap DY-549, SS-A488 or SS-A647 in AHL-/- for 15 minutes at room temperature. Brains were then mounted onto PLL coated coverslips and covered with AHL-/- for imaging. Imaging was performed with an LSM 880 scanning confocal microscope under a 20x objective water immersion objective (W Plan-Apochromat 20x/1.0 DIC Vis-IR M27 75mm). Z-stacks were taken under the following settings for 3 µm z-step size, 1024 x 1024 pixels, 0.59 µm x 0.59 µm pixel size, and 1.37 µs pixel dwell time. For deeper samples, the laser power was increased gradually as the focal plane depth increased to reduce the effect of light scattering.

**Table S3-3.** Confocal microscopy settings for brain registration

| Dye | Excitation | Emission |
|---|---|---|
| SS-A488 | 488 nm | 490-650 nm |
| SS-DY549 | 561 nm | 564 –739 nm |
| SS-A647 | 633 nm | 638-755 nm |

*Dye loading and imaging for registered brains*

Flies aged 10 days post eclosion were dissected in ice-cold AHL-/-. Brains were stored on ice in AHL-/- during the remainder of the dissections (no longer than 20 minutes) before being loaded for 15 minutes with 10 µM Surface Snap A647 in AHL-/- containing 0.2% Pluronic F127 (resuspended in DMSO). Brains were then mounted onto PLL coated coverslips and covered with AHL-/- for imaging. Imaging was performed with an LSM 880 scanning confocal microscope under a 20x objective water immersion objective (Type). Z-stacks were taken under the following settings for 1 µm z-step size, 1024x 1024 pixels, 0.59 x0.59 µm pixel size, and 1.03 µs pixel dwell time . For deeper samples, the laser power was increased gradually as the focal plane depth increased to reduce the effect of light scattering.

**Table S3-4.** Confocal microscopy settings for brain registration

| Dye | Excitation | Emission |
|---|---|---|
| SS-A647 | 633 nm | 638-755 nm |

*Image analysis.*

Each confocal stack was collapsed into a maximum intensity projection (Gal4 expression patterns) or average intensity projection (LexA>CD8::GFP expression patterns). For segmented images, projections were traced and filled using Simple Neurite Tracer (FIJI), and all regions not in that projection fill were removed from the image. This was performed to remove autofluorescence background in LexA>CD8::GFP flies which can be high in regions such as the optic lobes.

**Image analysis.**

Each confocal stack was collapsed into a maximum intensity projection (Gal4 expression patterns) or average intensity projection (LexA>CD8::GFP expression patterns). For segmented images, projections were traced and filled using Simple Neurite Tracer (FIJI), and all regions not in that projection fill were removed from the image. This was performed to remove autofluorescence background in LexA>CD8::GFP flies which can be high in regions such as the optic lobes.

**Registration.**

Confocal stacks were registered using the method described in Cachero et al. 2010[3]. Briefly, confocal stacks were reoriented manually in FIJI to best match template orientation and resized to 0.62µm x 0.62µm x 1µm pixel size. Brains were then registered to JFRC2010 using the FIJI CMTK Registration GUI[3] run directly through terminal. Quality was then assessed manually by assuring that the brain structures were not warped or oddly shaped due to the reformatting. We also overlaid the template brain and the registered brain to assure that brain structures were maintained in shape and structure. Finally, each brain was then cropped at a distinct anatomical landmark where the mushroom bodies end in the brain, and the z- plane was resliced so that each stack was 116 individual images. This final step was required to assure good alignment in the z-plane as only the first half of the brain accurately registers using our technique.

**Dice Coefficient Analysis.**

A Dice Coefficient defined areal overlaps between our registered brains and the template brains. Each region was then compared to the Ito region from that z-plane slice, and the Dice Coefficient was calculated using the following equation.

$$\mathrm{Dice}(R_z)=2\frac{|R_z^T \cap R_z^D|}{|R_z^T|+|R_z^D|}$$

Where is the area of the Ito $R_z^T$ ROI and $R_z^D$ is the area of the experimenter drawn ROI.

**Neuron fill generation.**

Registered LexA>CD8::GFP expressing lines were traced and filled using Simple Neurite Tracer.

**Symmetric Euclidean Distance.**

Symmetric Euclidean Distance was measured using the ROIs drawn for Dice Coefficient and the original Ito ROIs. The Symmetric Euclidean Distance was defined as:

$$\text{Symmetric Euclidean Distance}(R_z) = \frac{d_z^{D\text{-}T} + d_z^{T\text{-}D}}{2}$$

Where $d_z^{D\text{-}T}$ is the shortest distance from one point on the raw data to any point on the template data, and $d_z^{T\text{-}D}$ is the shortest distance from one point on the template data to any point on the raw data set.

**NBLAST search.**

NBLAST search was queried against 12,000 neurons of the FlyCircuit[5] database and all of the Janelia Gal4[6] database. The resulting query list contained approximately 15,500 neurons and expression patterns. We seeded this data base with a trace of the interoceptive neurons from immunohistochemistry. The target consisted of a single traced neurite (Simple Neurite Tracer, FIJI) from a live brain registered to JFRC2010. The seed and live brain ISN data were transformed from JFRC2010 template space to FCWB via a bridging registration. NBLAST similarity search was then performed using mean normalized comparisons, which compares the neurons bi-directionally to ensure a good match and alpha factor enabled, which weights long range projections more heavily. Neurons selected as positive hits by this search were defined as those with a positive mean normalized similarity score.

**Supporting Figures**

**Figure S1.** Alexa fluor dye loading in SNAP$_f$-CD4 transfected HEK293T cells

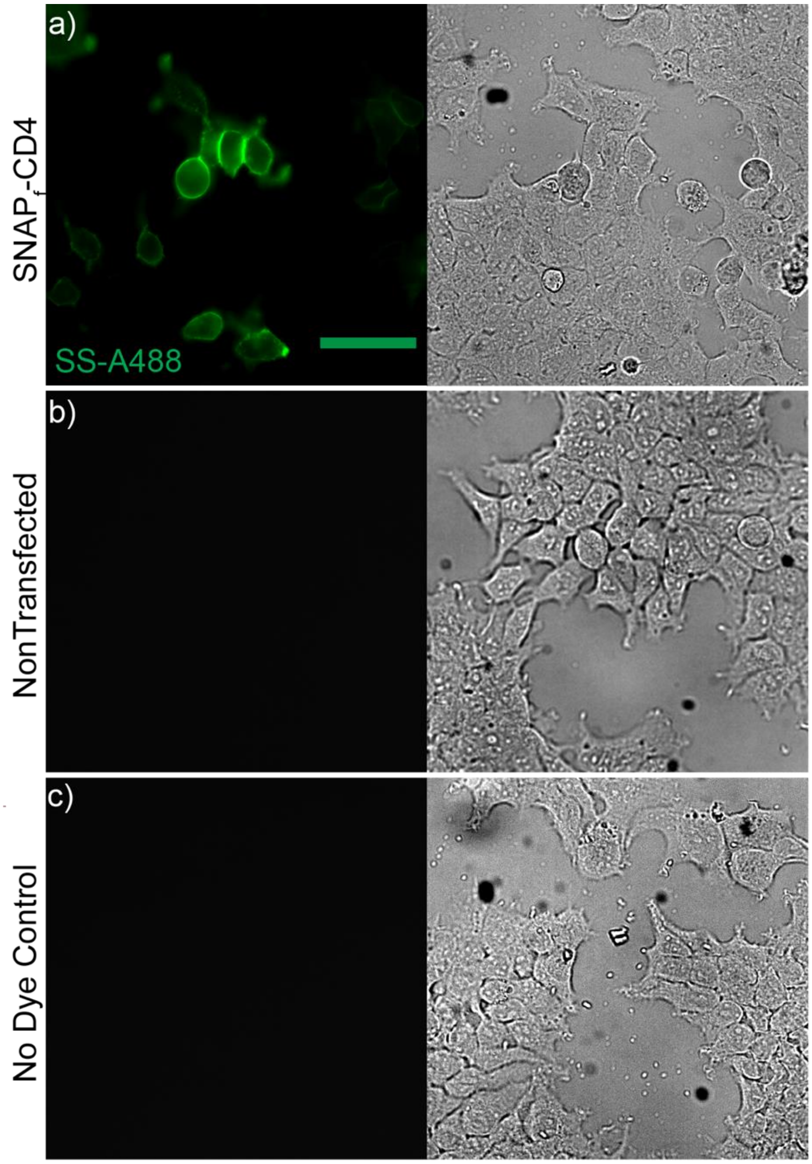

Alexa fluor dye loading in HEK293T cells expressing CMV promoted extracellular SNAP$_f$-CD4. Epifluorescence image of live cell dye loading of HEK293T cells expressing SNAP$_f$-CD4 (CMV promotor) and stained with **a)** SS-A488 (100 nm, green), with a DIC image of the same area (right). The white line depicts non-transfected cell's location in both DIC and live cell epifluorescence imaging. Controls were treated with **b)** dye but no transient transfection with SNAP$_f$-CD4 or **c)** transient transfection with SNAP$_f$-CD4 but no dye loading. Scale on all images is 50 µm.

**Figure S2.** Immunocytochemistry (ICC) in HEK293T cells

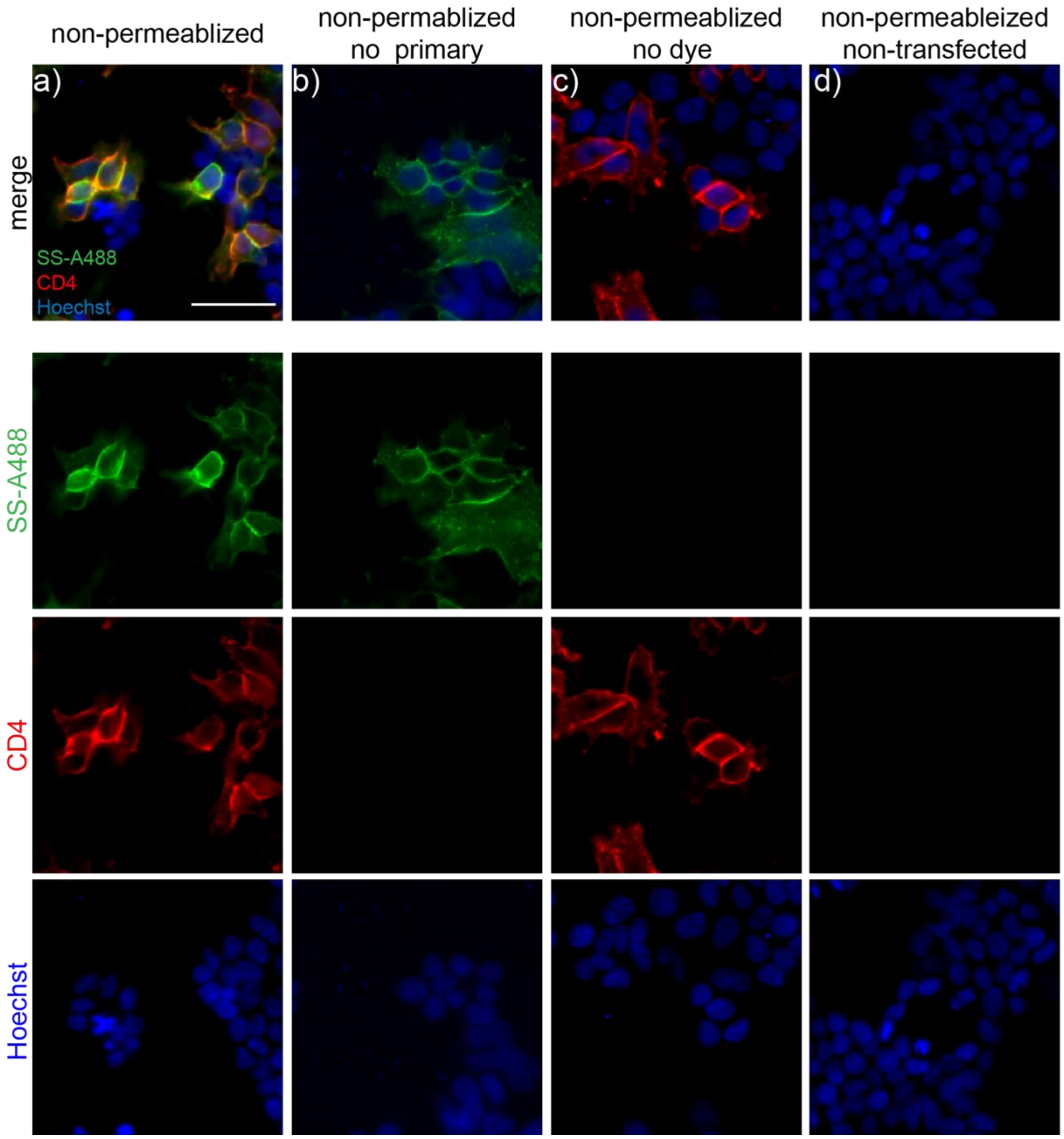

Epifluorescence images of HEK293T cells expressing SNAP$_f$-CD4 (CMV promotor) and stained with SS-A488 (100 nm, green as in Figure B). Cells were then fixed and stained under non-permeabilizing conditions for CD4 (red) and nuclear counterstained using Hoechst 33342 (at a concentration of 10 µg/ul, equivalent to 16 µM) **(a).** Controls were treated with **b)** dye but no primary CD4 antibody, **c)** primary antibody but no dye and d) with dye and primary antibody but without transfection. Scale bar is 50 µm.

**Figure S3.** Alexa fluor dye loading in HEK293TCells

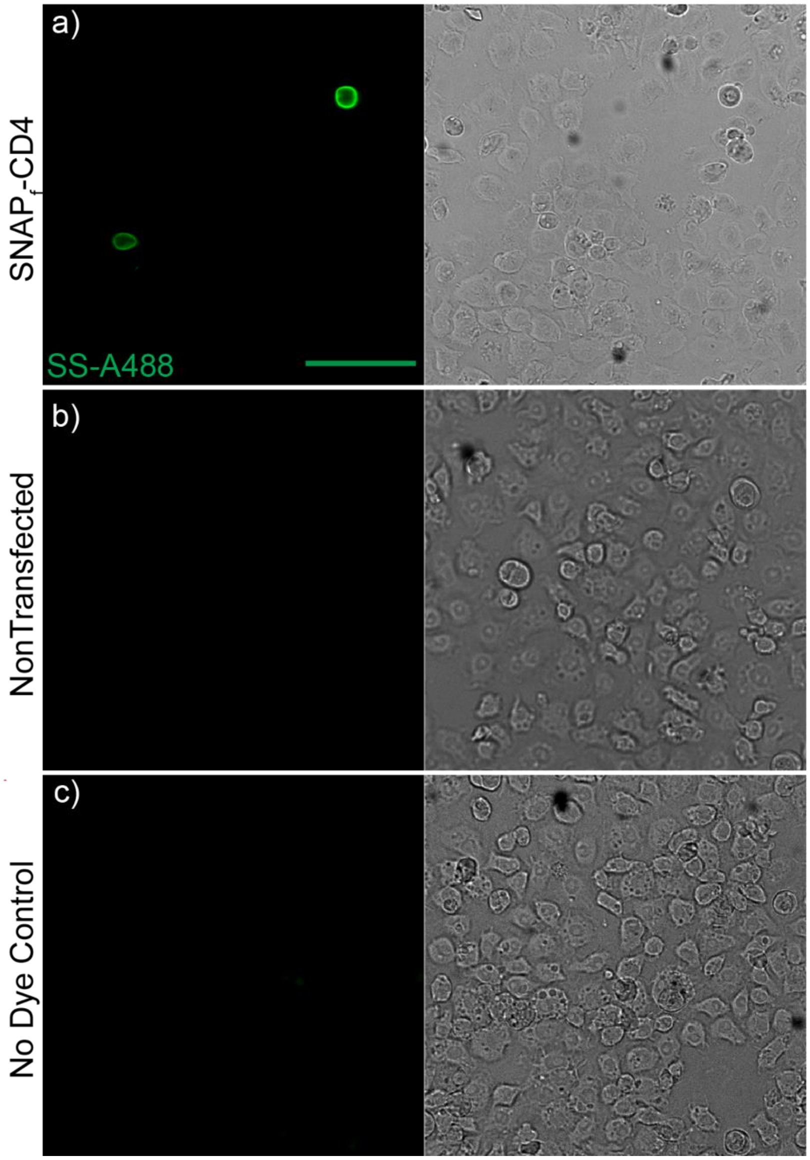

Alexa fluor dye loading in S2 cells expressing extracellular UAS-SNAP$_f$-CD4 under cotransfected pTubulin-Gal4. Epifluorescence image of live cell dye loading with SS-A488 (100 nm, green) in S2 cells expressing UAS-SNAP$_f$-CD4 after cotransfection with pTubulin-Gal4 (**a)**, A DIC image of the same area is to the right. The white line depicts non-transfected cell's location in both DIC and live cell epifluorescence imaging. Controls were treated Controls were treated with **b)** SS-A488 dye but no transient transfection with SNAP$_f$-CD4 or **c)** transient transfection with SNAP$_f$-CD4 but no dye addition. Scale on all images is 50 μm.

**Figure S4.** Immunocytochemistry in Drosophila S2 cells expressing SNAPf-CD4.

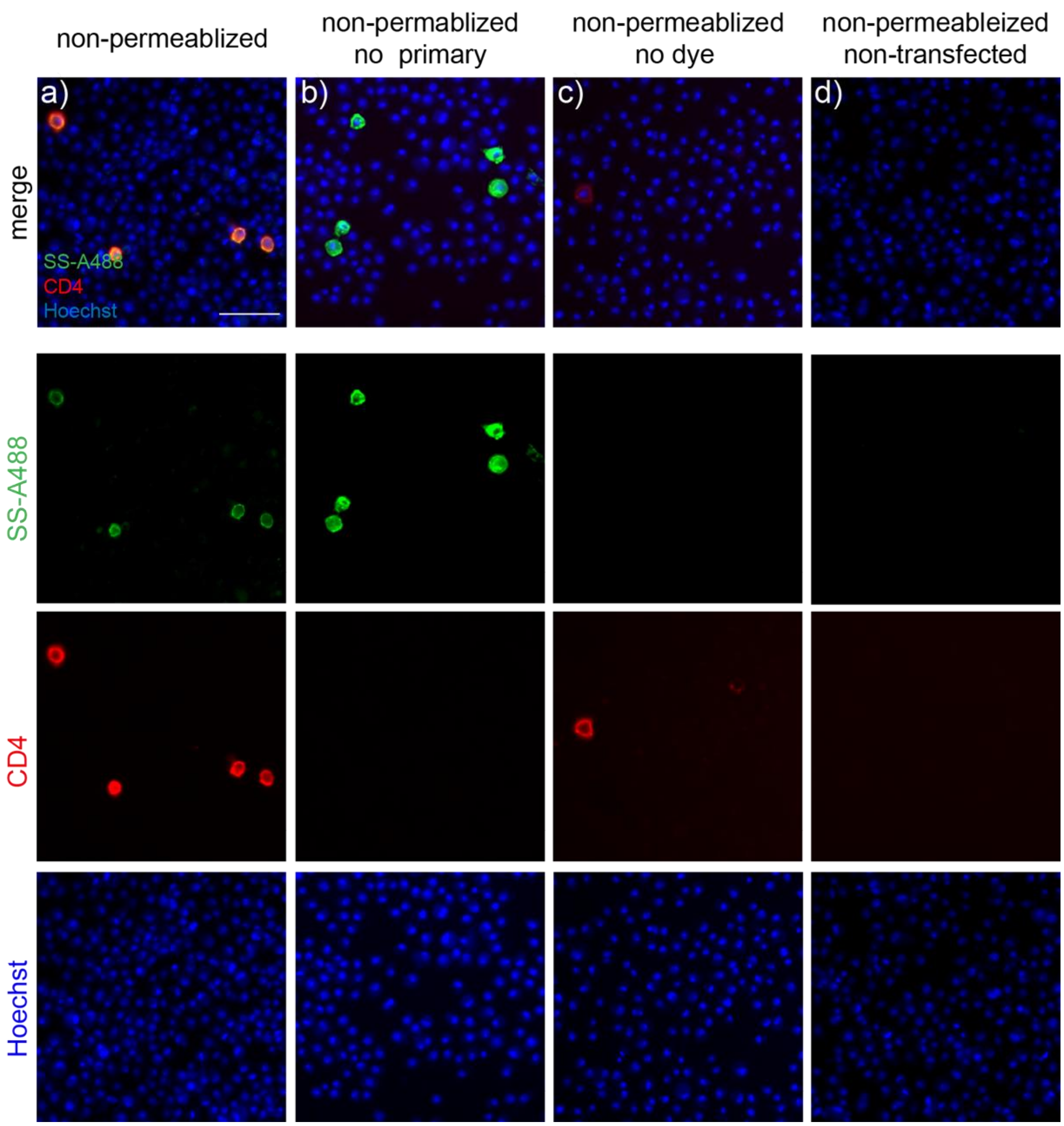

Confocal images of post hoc immunocytochemistry of *Drosophila* S2 cells expressing SNAP$_f$-CD4 (cotransfection with driver pTubulin Gal4 and UAS-SNAP$_f$-CD4) stained with SS-A488 (100 nM as in Figure C) fixed and stained for CD4 (red) under non-permeabilizing conditions **(a)**. Cells were nuclear counterstained using Hoechst 33342 (at a concentration of 10 µg/ul, equivalent to 16µM). Controls were treated with **b)** dye but no primary CD4 antibody, **c)** primary antibody but no dye and **d)** with dye and primary antibody but without transfection. Scale bar is 50 µm.

**Figure S5.** Immunohistochemistry for $SNAP_f$-CD4

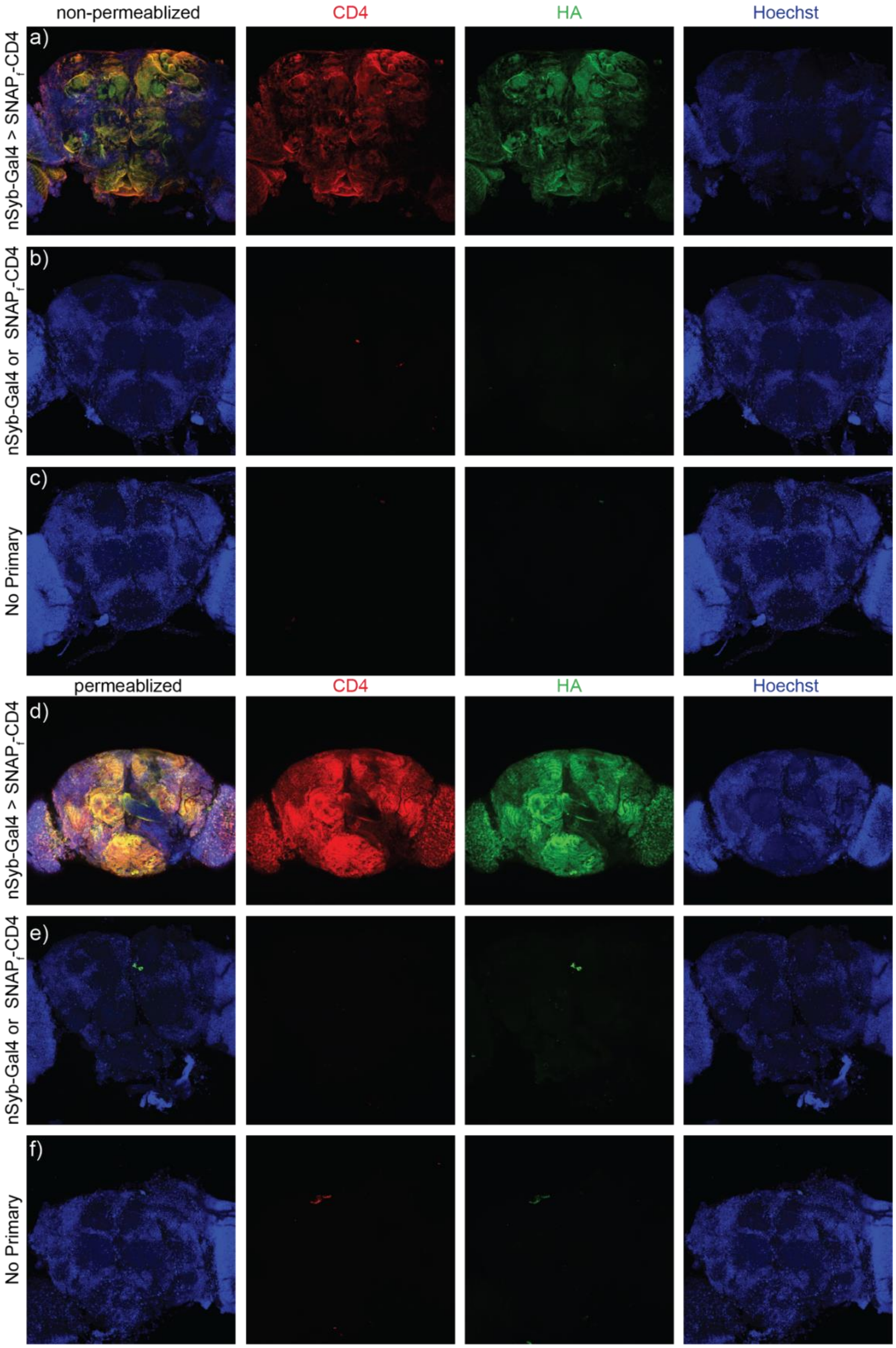

Permeabilized and non-permeabilized immunohistochemistry for $SNAP_f$-CD4 in panneuronal expressing nSyb-Gal4>$SNAP_f$-CD4 *Drosophila* brains. **a)** Maximum confocal z-projection of a fixed nSyb-Gal4, $SNAP_f$-CD4 brain stained under non-permeabilizing conditions for HA (green), CD4 (red) and nuclear counterstained using Hoechst 33342 (at a concentration of 10 µg/ul, equivalent to 16µM). Controls were treated under non-permeabilized conditions in the **b)** absence of transgene expression ($SNAP_f$-CD4 or nSyb-Gal4) or **c)** absence of primary antibody. **d)** Maximum confocal projection of nSyb-Gal4, $SNAP_f$-CD4 brain treated with triton x-100 prior to immunostaining for HA (green), CD4 (red) and nuclear counterstaining using Hoechst 33342. Controls were treated under permeabilizing conditions in the **e)** absence of transgene expression ($SNAP_f$-CD4 or nSyb-Gal4) or **f)** absence of primary antibody. Scale is 50 μm.

**Figure S6.** Dye loading in panneuronal LexA lines

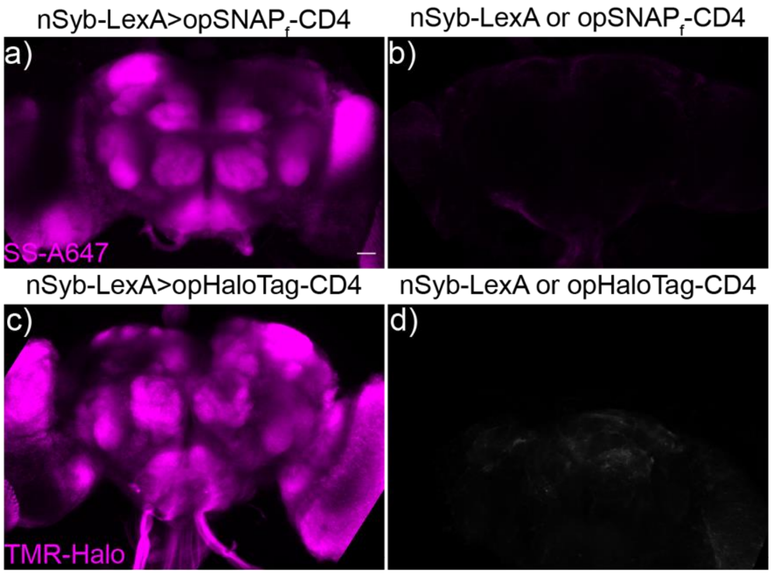

Dye loading in panneuronal op-SNAP$_f$-CD4 and op-HaloTag-CD4 LexA lines. Average confocal z- projection of **a)** nSyb-LexA>op-SNAP$_f$-CD4 brain or **b)** nSyb-LexA or op-SNAPf-CD4 genetic control brain loaded with SS-A647(10 µM). Confocal average z-projection of **c)** nSyb-LexA>op-HaloTag-CD4 brain or **d)** nSyb-LexA or op-HaloTag-CD4 brain loaded with TMR-Halo (10 µM). Scale for all images is 50 µm.

**Figure S7.** Intracellular vs. extracellular genetic targeting of dye molecules

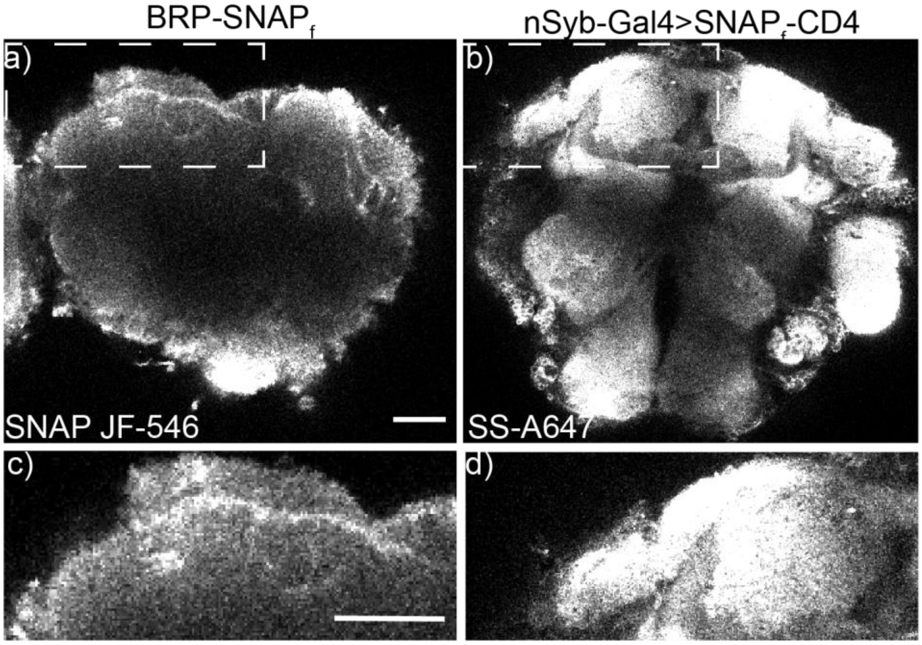

Genetic targeting of dye molecules in *Drosophila* brain using intracellularly targeted BRP-SNAP$_f$ and extracellularly targeted SNAP$_f$-CD4. **a)** Single plane confocal image of brain expressing BRP-SNAP$_f$ on the intracellular surface loaded with SNAP JF-546 cell permeant dye. White box denotes region of protocerebrum selected for zoomed in inlay in panel **c). b)** Single plane confocal image of brain expressing panneuronal SNAP$_f$-CD4 on the extracellular surface loaded with SS-A647 (10 μM). **d)** zoomed in region of protocerebrum from panel **b)** (denoted by white box). Scale for all images is 50 μm.

**Figure S8.** Dice Coefficient across z planes

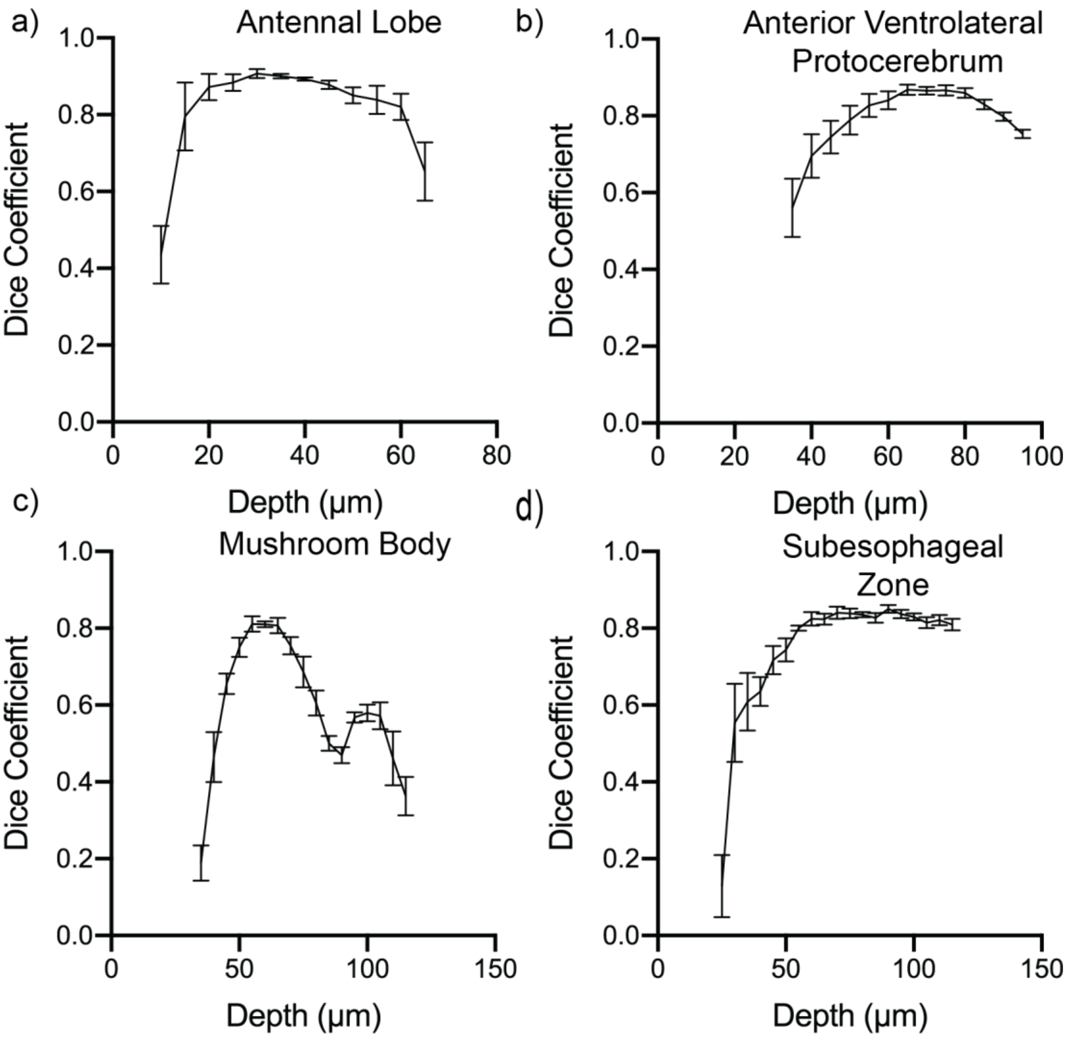

Dice coefficient across z planes at 5 µm steps throughout each anatomical structure. Average dice coefficient ± SEM for each individual plane taken at 5 µm steps through the first half of the brain or to the end of the structure for **a)** Antennal Lobe, **b)** Anterior Ventrolateral Protocerebrum, **c)** Mushroom Body, or **d)** Suboesophageal Zone. Data represent the average Dice coefficient ± SEM across 10 individual registered brains (5 male and 5 female).

**Figure S9.** Symmetric Euclidean Distance across z-planes

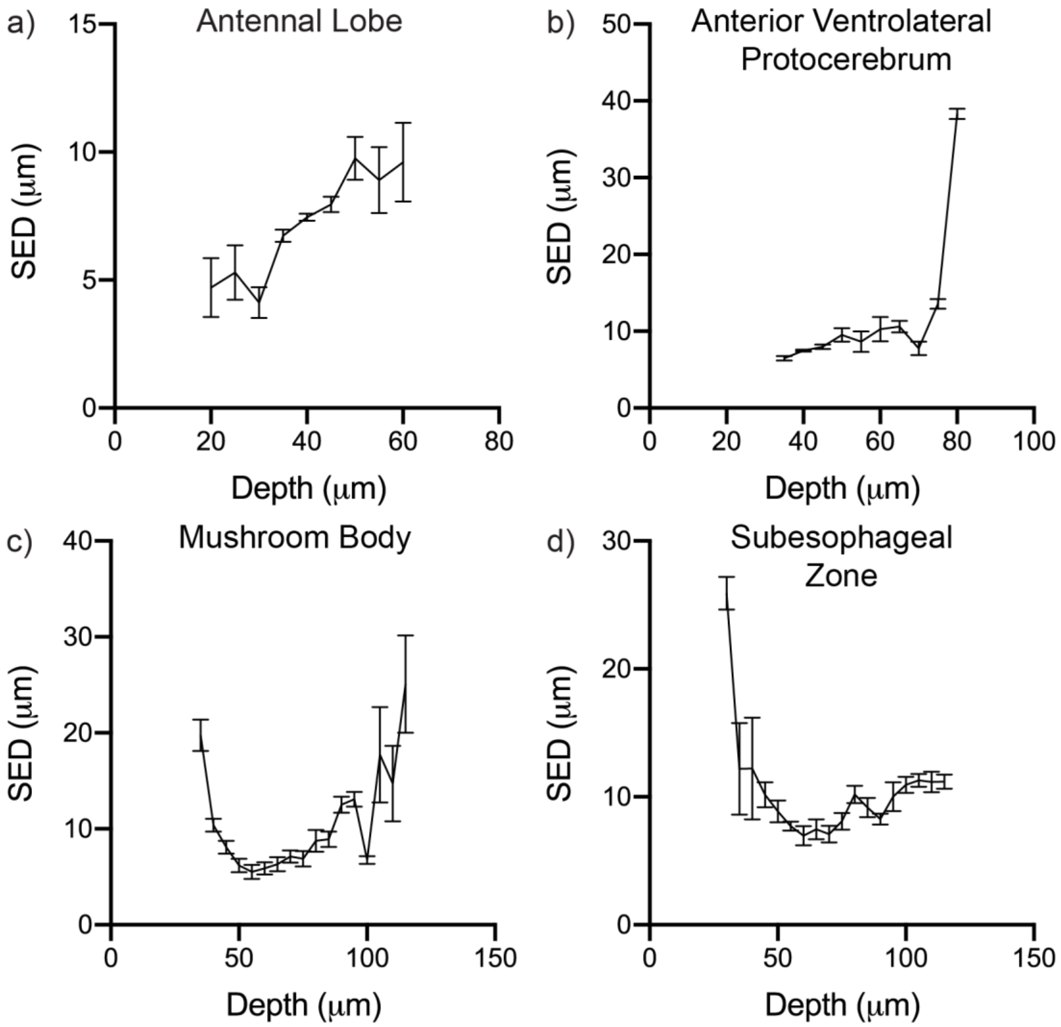

Average Symmetric Euclidean Distance across z planes at 5 μm steps throughout each anatomical structure. Average Symmetric Euclidean Distance across z planes ± SEM for each individual plane taken at 5 μm steps through the first half of the brain or to the end of the structure for **a)** Antennal Lobe, **b)** Anterior Ventrolateral Protocerebrum, **c)** Mushroom body, or **d)** Suboesophageal Zone. Data represent the average Symmetric Euclidean Distance for that z plane ± SEM across 10 individual registered brains (5 male and 5 female).

***References.***
